# A novel *RLIM/RNF12* variant disrupts protein stability and function to cause severe Tonne-Kalscheuer syndrome

**DOI:** 10.1101/2020.12.09.417873

**Authors:** Francisco Bustos, Carmen Espejo-Serrano, Anna Segarra-Fas, Alison J. Eaton, Kristin D. Kernohan, Meredith J. Wilson, Lisa G. Riley, Greg M. Findlay

**Affiliations:** MRC Protein Phosphorylation & Ubiquitylation Unit, University of Dundee, Dundee, UK; Department of Medical Genetics, University of Alberta, Edmonton, Alberta, Canada; Newborn Screening Ontario, Children’s Hospital of Eastern Ontario, Ottawa, Canada; Children’s Hospital of Eastern Ontario Research Institute, University of Ottawa, Ottawa, Canada; Department of Clinical Genetics, The Children’s Hospital at Westmead, Sydney, Australia; Discipline of Genomic Medicine, University of Sydney, Sydney, Australia; Rare Diseases Functional Genomics, Kids Research, The Children’s Hospital at Westmead and The Children’s Medical Research Institute, Sydney, Australia; Discipline of Child & Adolescent Health, Sydney Medical School, University of Sydney, Sydney, Australia

## Abstract

Tonne-Kalscheuer syndrome (TOKAS) is an X-linked intellectual disability syndrome associated with variable clinical features including craniofacial abnormalities, hypogenitalism and diaphragmatic hernia. TOKAS is caused exclusively by variants in the gene encoding the E3 ubiquitin ligase gene *RLIM*, also known as. Here we report identification of a novel *RLIM* missense variant, c.1262A>G p.(Tyr421Cys) adjacent to the regulatory basic region, which causes a severe form of TOKAS resulting in perinatal lethality by diaphragmatic hernia. Inheritance and X-chromosome inactivation patterns implicate *RLIM* p.(Tyr421Cys) as the likely pathogenic variant in the affected individual and within the kindred. We show that the RLIM p.(Tyr421Cys) variant disrupts both expression and function of the protein in an embryonic stem cell model. RLIM p.(Tyr421Cys) is correctly localised to the nucleus, but is readily degraded by the proteasome. The RLIM p.(Tyr421Cys) variant also displays significantly impaired E3 ubiquitin ligase activity, which interferes with RLIM function in *Xist* long-non-coding RNA induction that initiates imprinted X-chromosome inactivation. Our data uncover a highly disruptive missense variant in *RLIM* that causes a severe form of TOKAS, thereby expanding our understanding of the molecular and phenotypic spectrum of disease severity.

## INTRODUCTION

Tonne-Kalscheuer syndrome (TOKAS; MIM #300978) is a recently described X-linked recessive multiple congenital anomaly disorder^1-3^. Male patients display global developmental delay apparent from early infancy, impaired intellectual development, speech delay, behavioural abnormalities, and abnormal gait. Affected individuals also display dysmorphic facial features, anomalies of the hands, feet and nails, abnormal pulmonary development, and urogenital abnormalities with hypogenitalism. In a subset of severely affected patients, development of congenital diaphragmatic hernia *in utero* may result in perinatal or premature death. Carrier females may display minor clinical manifestations including very mild skeletal or hormonal abnormalities^1^.

In all reported cases, TOKAS is caused by variants in the RING finger type E3 ubiquitin ligase *RLIM/RNF12*, which ubiquitylates transcription factor substrates to control key developmental processes including imprinted X-chromosome inactivation^4^, stem cell maintenance and differentiation^5,6^. In all cases, patient-derived RLIM TOKAS variants lead to impaired E3 ubiquitin ligase activity^1,6^ without major impact on other aspects of RLIM expression and function, including protein stability, phosphorylation, subcellular localisation and protein:protein interactions^6^. Specific disruption of RLIM activity by TOKAS variants result in deregulated stem cell differentiation to neurons^6^, providing insight into the cellular processes that may underpin TOKAS etiology.

Thus far, 8 variants that cause TOKAS in distinct unrelated families have been reported^1-3^. *RLIM* variants have been reported in 4 cases out of a cohort of 405 cases of unresolved syndromic X-linked intellectual disability with no known genetic or environmental basis^2^. This data suggests that a significant proportion of unresolved X-linked intellectual disability cases may be caused by *RLIM* variants, and that other *RLIM* TOKAS variants await identification.

Here, we report identification of a novel *RLIM* missense variant, p.(Tyr421Cys), adjacent to the regulatory basic region, which causes a severe form of TOKAS leading to perinatal lethality by diaphragmatic hernia. Inheritance and X-chromosome inactivation patterns clearly implicate *RLIM* p.(Tyr421Cys) as the causative variant in the affected individual and kindred. We show that the RLIM p.(Tyr421Cys) variant severely disrupts protein expression and function, and is readily degraded by the proteasome. RLIM p.(Tyr421Cys) also displays significantly impaired E3 ubiquitin ligase activity. Together, defects in RLIM p.(Tyr421Cys) protein expression and activity profoundly interfere with RLIM function in *Xist* long-non-coding RNA induction, a key step in initiating imprinted X-chromosome inactivation. Our data uncover a highly disruptive missense variant in *RLIM* that causes a severe form TOKAS, thereby expanding our understanding of the molecular and phenotypic spectrum of disease severity.

## RESULTS

### Clinical information for a patient with an undiagnosed developmental disorder

The male proband was the first child of unrelated parents. The first trimester nuchal translucency/morphology scan and second trimester fetal morphology scan were reported as normal. Polyhydramnios was detected at 36 weeks’ gestation and intrauterine growth restriction (IUGR) was reported at 37 weeks. Labour was induced at 39 weeks and delivery was via emergency Caesarean section for subsequent fetal distress. The baby became cyanosed immediately after birth, was unable to be resuscitated and died at 30 minutes of age.

On examination the baby was symmetrically small for gestational age with a birth weight of 2285 g (<3^rd^ percentile), length 48.5 cm (10^th^ percentile) and head circumference 32.5cm (<3^rd^ percentile). Craniofacial anomalies were identified, including hypertelorism, broad nasal bridge with flat nasal tip, and a high arched palate. He had distal limb hypoplasia, with brachytelephalangy, soft tissue syndactyly between the 2^nd^ and 3^rd^ toes bilaterally and absent or hypoplastic nails on 2^nd^-3^rd^ fingers and 2^nd^-4^th^ toes. There were no ocular abnormalities.

Post mortem examination revealed congenital diaphragmatic hernia with aplasia of the posterolateral left hemi-diaphragm, displacement of the mediastinum to the right, herniation of the small and large intestine, stomach, spleen, left lobe of liver, and pancreas into the left hemithorax, absent middle lobe right lung and severe bilateral pulmonary hypoplasia. There was an accessory spleen. The cardiovascular system showed aberrant aortic arch branching with the left subclavian, left common carotid and right common carotid arteries arising from a single brachiocephalic trunk. Urogenital abnormalities included cryptorchidism with pelvic testes, but kidneys were normal. A cavum septum pellucidum was present. The clinical features were considered to be most consistent with Fryns syndrome (MIM #229850). Pallister-Killian syndrome (tetrasomy 12p; MIM #601803) was considered, but fluorescence in-situ hybridisation for chromosome 12p on fetal lung imprints, standard karyotype on cultured fibroblasts and chromosome microarray on DNA extracted from stored fetal tissue (Agilent Sureprint G3 ISCA Targeted Microarray 8×60K) were normal.

### Identification of a novel missense *RLIM* variant and diagnosis of Tonne-Kalscheuer syndrome (TOKAS)

In order to investigate the genetic basis of this case, we performed whole exome sequencing on genomic DNA from the proband and his parents, and identified a maternally inherited hemizygous variant in the X-linked *RLIM* gene (NM_016120.3) c.1262A>G, p.(Tyr421Cys) (Figure 1A). *RLIM* encodes the RLIM/RNF12 E3 ubiquitin ligase that is mutated in the recessive X-linked disorder Tonne-Kalscheuer syndrome (TOKAS)^1-3^, which is characterised by clinical features that significantly overlap with Fryns syndrome. There were no variants in *PIGN*, in which biallelic loss of function mutations have been reported to cause a Fryns-like syndrome. We hypothesised that the individual was affected by a severe form of TOKAS that is caused by this novel *RLIM* variant. Consistent with this notion, the p.(Tyr421Cys) variant is located adjacent to the basic region of the RLIM protein, a key regulatory domain that is mutated in several TOKAS kindreds^1-3^. *RLIM* p.(Tyr421Cys) has not been observed in affected individuals in the literature and was not reported in the ClinVar archive of genomic variation in human health (https://www.ncbi.nlm.nih.gov/clinvar) or the gnomAD population database (https://gnomad.broadinstitute.org/). As a result, the variant was submitted to ClinVar under the accession number SCV001435291.

**Figure 1:**
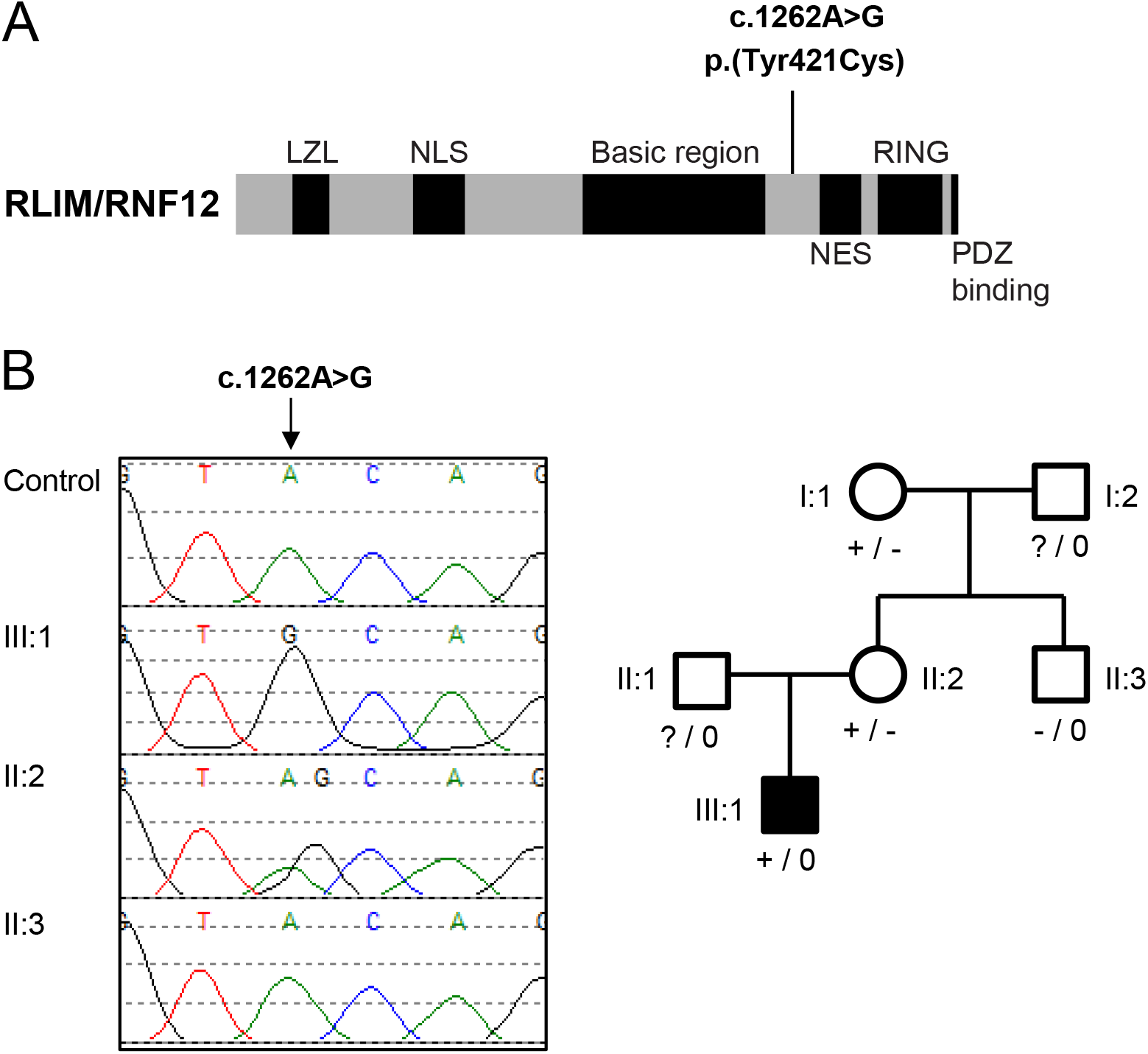
Identification of RLIM p.(Tyr421Cys), a novel variant in a severe form of Tonne-Kalscheuer Syndrome. A) Schematic diagram of the RLIM protein domain structure, with the position of the c.1262A>G p.(Tyr421Cys) variant identified by exome sequencing indicated. B) (Left panel) Genomic DNA sequencing electropherogram identifying the *RLIM* c.1262A>G p.(Tyr421Cys) variant within the kindred. (Right panel) Family pedigree showing inheritance of the *RLIM* c.1262A>G p.(Tyr421Cys) variant.

Sanger sequencing confirmed the presence of the *RLIM* c.1262A>G p.(Tyr421Cys) variant in the proband, whilst the mother and maternal grandmother were confirmed as carriers (Figure 1B). The variant was absent in the mother’s unaffected brother (Figure 1B), consistent with an X-linked inheritance pattern. Furthermore, X-chromosome inactivation analysis showed the mother had a highly skewed X-chromosome inactivation pattern (95%; Supplementary Figure 1), which is characteristic of female carriers of *RLIM* TOKAS variants^1,3^. The clinical features and identification of this novel *RLIM* variant inherited by the affected individual led to a revised diagnosis of a severe form of TOKAS.

### RLIM p.(Tyr421Cys) is poorly expressed and readily degraded by the proteasome

The diagnosis of severe TOKAS prompts the hypothesis that the p.(Tyr421Cys) variant impacts on RLIM protein expression and/or function. This variant is predicted to be damaging to the protein by multiple *in silico* programs (SIFT (v6.20)^7^: deleterious (score: 0); PolyPhen2^8^: probably damaging (score: 0.994); MutationTaster (v2013)^9^: disease causing (p-value: 1)), and is classified as a variant of uncertain significance (VUS) by ACMG guidelines^10^. Therefore, we exploited RLIM-deficient male mouse Embryonic Stem Cells (mESCs) to investigate expression and function of RLIM p.(Tyr421Cys) in a cellular model. In comparison to wild-type human RLIM, p.(Tyr421Cys) RLIM protein is poorly expressed (Figure 2A). Quantitative analysis indicates that RLIM p.(Tyr421Cys) expression is reduced to 28.7 ± 11.5 % of wild-type (Figure 2B). However, the mRNA is expressed at similar levels, (Figure 2C) and the protein is correctly localised to the nucleus (Figure 2D), suggesting that the RLIM p.(Tyr421Cys) variant specifically interferes with translation and/or stability. We explored whether RLIM p.(Tyr421Cys) is more readily turned over by the proteasome. Indeed, treatment of RLIM p.(Tyr421Cys) expressing mESCs with the proteasome inhibitor MG132 expression rescues RLIMp.(Tyr421Cys) expression to levels approaching that of wild-type RLIM (Figure 2E), indicating that RLIM p.(Tyr421Cys) more readily undergoes proteasomal degradation. Taken together, these data indicate that the RLIM p.(Tyr421Cys) variant destabilises the protein.

**Figure 2:**
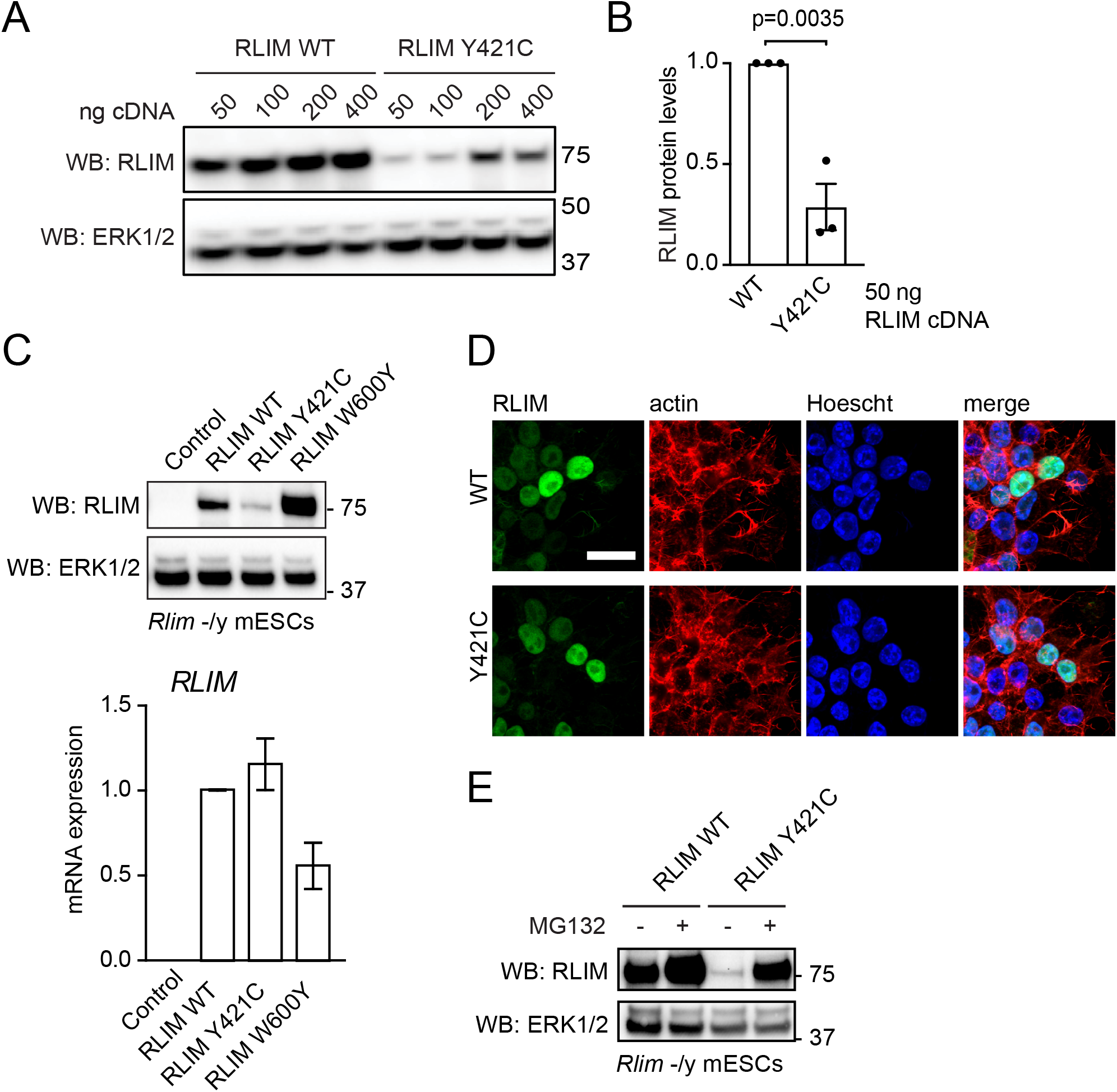
RLIM p.(Tyr421Cys) is poorly expressed and readily degraded by the proteasome. A) Human RLIM wild-type (WT) or Y421C TOKAS variant were expressed at increasing amounts (50-400 ng plasmid DNA) in *Rlim*^-/y^ mESCs. RLIM expression was determined by immunoblotting and ERK1/2 expression analysed as a loading control. B) Quantification of human RLIM wild-type (WT) or Y421C TOKAS variant protein expression from 50 ng plasmid cDNA in *Rlim*^-/y^ mESCs. RLIM expression was determined by immunoblotting. Normalised data are represented as mean ± standard error of the mean (n=3). Statistical significance was determined by student’s t-test. C) Human RLIM wild-type (WT), Y421C TOKAS variant or W600Y catalytically-inactive variant were expressed in *Rlim*^-/y^ mESCs. (Top panel) Human RLIM protein expression was determined by immunoblotting and ERK1/2 expression analysed as a loading control. (Bottom panel) Human *RLIM* mRNA expression was determined by qRT-PCR and normalised to *Gapdh* mRNA expression. Data are represented as mean ± standard deviation (n=2). D) Human RLIM wild-type (WT) and Y421C TOKAS variant were expressed in *Rlim*^-/y^ mESCs. RLIM localisation was determined by immunofluorescence. E) Human RLIM wild-type (WT) or Y421C TOKAS variant were expressed in *Rlim*^-/y^ mESCs treated with vehicle or 10 μM MG132 for 4 h. RLIM expression was determined by immunoblotting and ERK1/2 expression analysed as a loading control.

### RLIM p.(Tyr421Cys) variant interferes with E3 ubiquitin ligase activity

As other RLIM TOKAS variants located within the proximal basic region (Figure 1A) have been shown to disrupt E3 ubiquitin ligase activity^1,6^, we next sought to determine the impact of p.(Tyr421Cys) on RLIM catalytic activity. We expressed and purified RLIM wild-type and p.(Tyr421Cys) from *E*.*coli*, and examined the ability of these variants to transfer ubiquitin from a cognate E2 (UBE2D1) to the prototypic substrate REX1, which is a transcription factor that is ubiquitylated by RLIM to initiate X-chromosome inactivation^11^. Wild-type RLIM efficiently promotes REX1 ubiquitylation, as evidenced by appearance of ubiquitylated species of increasing molecular weight upon addition of substrate (Figure 3A). However, RLIM p.(Tyr421Cys) significantly impairs REX1 ubiquitylation activity (Figures 3A), suggesting that the p.(Tyr421Cys) variant also has a negative impact on RLIM catalytic activity. Quantitative analysis indicates that RLIM p.(Tyr421Cys) activity is reduced to 32 ± 3.7% of wild-type (Figure 3B). Therefore, similar to other previously reported RLIM TOKAS variants, p.(Tyr421Cys) displays impaired E3 ubiquitin ligase activity.

**Figure 3:**
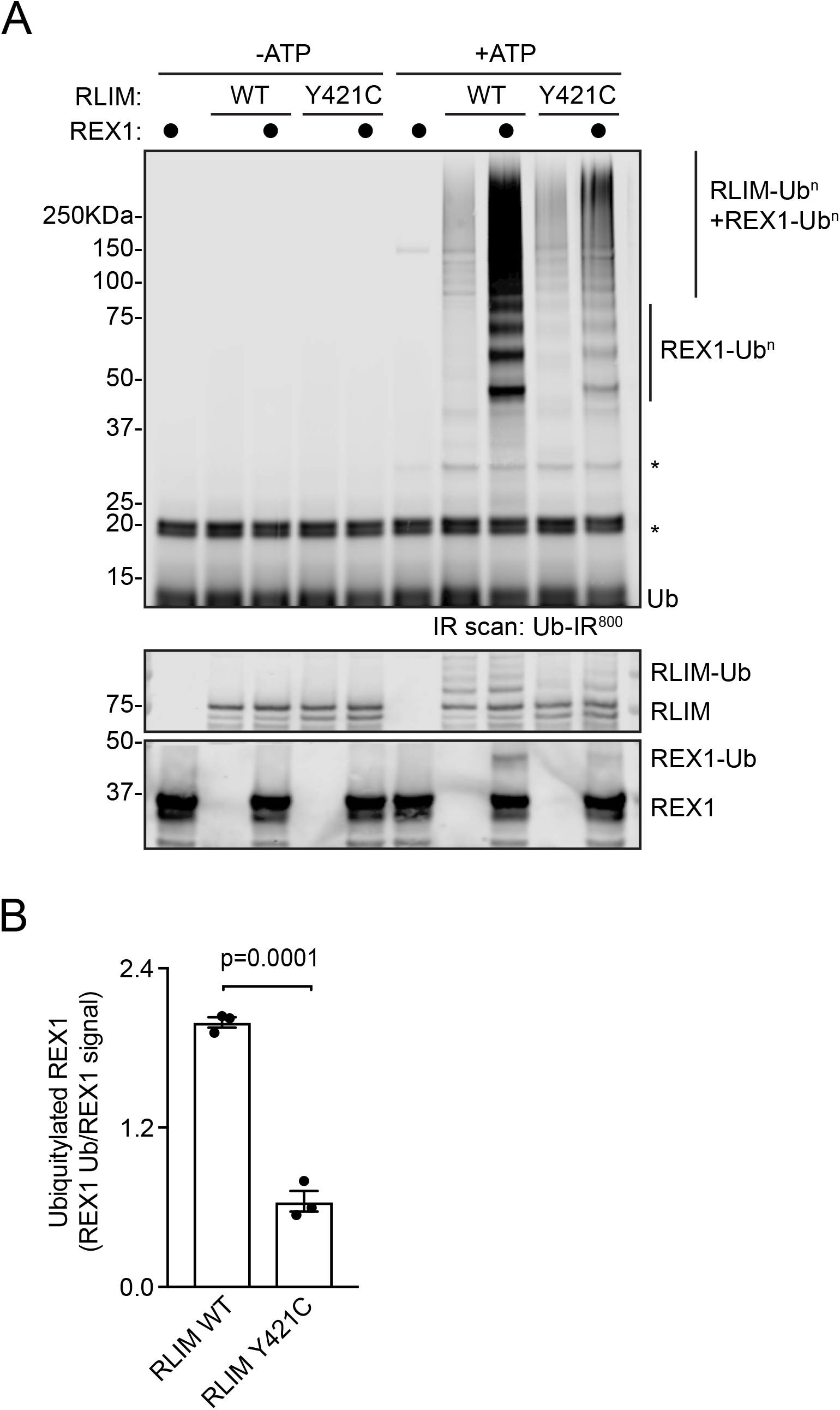
The p.(Tyr421Cys) variant impairs RLIM catalytic E3 ubiquitin ligase activity and substrate ubiquitylation. A) Recombinant human RLIM wild-type (WT) or Y421C TOKAS variant were assayed for E3 ubiquitin ligase activity by incubating with UBE1 E1, UBE2D1 E2 conjugating enzyme in the presence or absence of ATP and recombinant REX1 substrate. REX1-specific substrate ubiquitylation (REX1-Ub^n^) and RLIM, REX1 and/or free ubiquitin chains (RLIM -Ub^n^ + REX1-Ub^n^) are indicated. RLIM and REX1 protein levels were determined by immunoblotting, and ubiquitylated RLIM and REX1 indicated. B) Quantification of E3 ubiquitin ligase activity of RLIM wild-type (WT) or Y421C TOKAS variant. Data are represented as mean ± standard error of the mean (n=3). Statistical significance was determined by student’s t-test.

### Functional disruption of the RLIM signalling pathway by the p.(Tyr421Cys) variant

Finally, we explored the functional impact of the RLIM p.(Tyr421Cys) variant in a cellular model of RLIM signalling. RLIM plays a key role in imprinted X-chromosome inactivation^12^, which is initiated by RLIM -dependent transcriptional induction of the *Xist* long non-coding mRNA^13^. Thus, we used *Xist* expression as a readout for RLIM function. As expected, *Xist* expression in RLIM -expressing *Rlim*^+/y^ mESCs is very low (Figure 4A). However, *Xist* expression is induced by expression of wild-type human RLIM, but not by a catalytically inactive variant (W600Y; Figure 4A). When compared to wild-type RLIM, *Xist* induction is severely impaired upon expression of RLIM p.(Tyr421Cys) (Figure 4A), demonstrating that the p.(Tyr421Cys) variant profoundly interferes with RLIM function. In order to control for expression, we analysed RLIM levels in *Rlim*^-/y^ mESCs, which do not express endogenous RLIM (Figure 4B). Taken together, our data indicate that the RLIM p.(Tyr421Cys) variant disrupts RLIM stability and E3 ubiquitin ligase catalytic activity, which leads to significantly impaired RLIM function *in vivo*.

**Figure 4:**
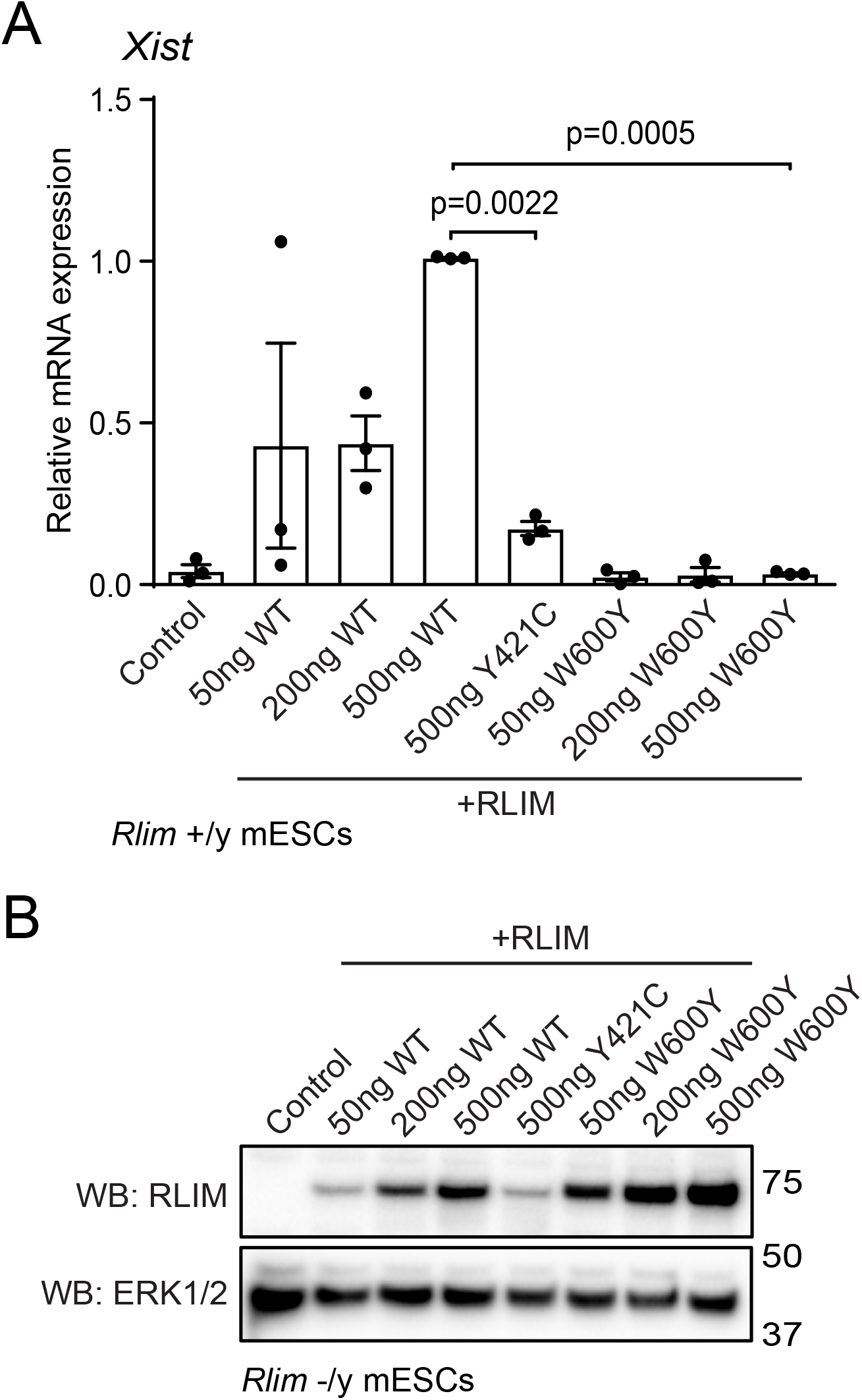
RLIM p.(Tyr421Cys) is functionally impaired in an assay for *Xist* lncRNA induction. A) Human RLIM wild-type (WT), Y421C TOKAS variant or W600Y catalytically-inactive variant were expressed at increasing amounts (50-500 ng plasmid DNA) in *Rlim*^+/y^ mESCs. *Xist* lncRNA expression was determined by qRT-PCR and normalised to *Gapdh* mRNA expression. Data are represented as mean ± standard error of the mean (n=3). Statistical significance was determined by one-way ANOVA. B) Human RLIM wild-type (WT), Y421C TOKAS variant or W600Y catalytically-inactive variant were expressed at increasing amounts (50-500 ng plasmid DNA) in *Rlim*^-/y^ mESCs. RLIM expression was determined by immunoblotting and ERK1/2 expression analysed as a loading control.

## DISCUSSION

Tonne-Kalscheuer syndrome (TOKAS) is a developmental disorder characterised by clinical features including intellectual disability, facial dysmorphism, velopharyngeal abnormalities and diaphragmatic hernia. In the most severe cases, diaphragmatic hernia causes death shortly after birth. TOKAS is caused by variants in the X-linked RLIM/RNF12 E3 ubiquitin ligase, which impair catalytic activity to varying extents. However, the extent to which phenotypic/disease severity correlates with genotypic severity (i.e. the extent of RLIM functional disruption) remains unclear.

Here, we provide a case report of a male patient who died shortly after birth with diaphragmatic hernia, facial dysmorphism and skeletal abnormalities. Cytogenetic studies showed no evidence of Pallister-Killian syndrome and an initial diagnosis of Fryns syndrome was thought most likely, although TOKAS was not considered as this baby was born several years before the association of diaphragmatic hernia with severe forms of TOKAS was delineated^1^. Research exome sequencing through the Care4Rare Canada Consortium uncovered a missense *RLIM* variant, p.(Tyr421Cys), which was heterozygous in the mother but not present in unaffected relatives, suggesting that the patient suffered a severe form of TOKAS. The mother showed a highly skewed pattern of X-chromosome inactivation, consistent with the presence of a highly deleterious X-linked variant. These findings raise the important question of whether other individuals diagnosed with Fryns syndrome and/or syndromic diaphragmatic hernia might actually represent incorrectly diagnosed TOKAS patients.

In this study, we explore the impact of the p.(Tyr421Cys) variant on RLIM protein expression and function. Strikingly, we find that RLIM p.(Tyr421Cys) is poorly expressed in an RLIM-deficient mouse embryonic stem cell model, and is prone to proteasomal degradation, although the protein is correctly localised in the nucleus. This is in contrast to other RLIM TOKAS variants, which show impaired catalytic activity but no major impact on stability^1,6^. Interestingly, analysis of the catalytic activity of recombinant RLIM p.Tyr421Cys indicates that this variant also displays impaired E3 ubiquitin ligase activity. This is consistent with a poorly understood function in catalysis of the basic region, which is proximal to the p.(Tyr421Cys) variant. Our data also support the notion that disrupted RLIM p.(Tyr421Cys) protein expression and catalytic activity together contribute to impairment of RLIM function. In this regard, induction of the *Xist* lncRNA by the RLIM p.(Tyr421Cys) variant, which is a key initiating step of X-chromosome inactivation, is significantly impaired.

In summary, we introduce RLIM p.(Tyr421Cys) as the prototypic member of a new class of RLIM TOKAS variant that impacts on both protein expression/stability and catalytic activity, which leads to severe TOKAS. This in turn expands our understanding of the molecular and phenotypic spectrum of TOKAS syndrome severity.

## METHODS

### Genomic DNA sequencing and analysis

Trio exome sequencing of genomic DNA from proband and parents was performed through a collaboration with Care4Rare program (http://care4rare.ca). Target capture was performed with the Agilent CRE V1.0 and sequencing performed on the Illumina NextSeq 500 using 150 base-pair paired-end reads. Data analysis was done by standard methods and variants annotated using both Annovar and custom scripts to identify whether they affect protein coding sequence, and whether previously seen in dbSNP132, the 100 Genomes dataset (Nov 2011), the NHLBI GO exomes or in the approx. 1500 exomes previously sequenced at the center^14^. Variant reporting relevant to diagnosis was informed by reference to HGMD, dbSNP, online search engines e.g. PubMed and locus-specific databases. Variants seen in >20 of their controls, or with an allele frequency of >3% in 100 Genomes or NHBLI were removed. Confirmatory Sanger sequencing and familial testing was performed in the Sydney Genomic Diagnostic laboratory, Children’s Hospital at Westmead.

### Mouse embryonic stem cell (mESC) culture

*Rlim*^+/y^ and *Rlim*^-/y^ mESCs were described in^1,6^. Cells were cultured on 0.1% gelatin (w/v)-coated plates in DMEM containing 10% (v/v) fetal bovine serum (FBS), 5% (v/v) Knock-Out serum replacement, 20 ng/ml GST-tagged leukemia inhibitory factor (LIF), penicillin/streptomycin, 2 mM glutamine, 0.1 mM minimum essential media (MEM) Non-essential amino acids, 1 mM sodium pyruvate (all Thermo Fisher Scientific) and 0.1 mM β-mercaptoethanol (Sigma-Aldrich) in a controlled atmosphere at 5% CO2 and 37°C.

### cDNA expression vectors and transfection

mESCs were transfected with Lipofectamine LTX (Thermo Fisher Scientific) according to manufacturer instructions. Plasmids used were pCAGGS human RLIM (DU53765), human RLIM Y421C (DU61099) and RLIM W600Y (DU53985). All cDNA clones were generated by MRC-PPU Reagents & Services; see http://mrcppureagents.dundee.ac.uk for detailed information and plasmid requests.

### Immunoblotting

Cells were harvested in lysis buffer containing 20 mM Tris (pH 7.4), 150 mM NaCl, 1 mM EDTA, 1% Nonidet P-40 (NP-40) (v/v), 0.5% sodium deoxycholate (w/v), 10 mM β-glycerophosphate, 10 mM sodium pyrophosphate, 1 mM NaF, 2 mM Na_3_VO_4_, and Roche Complete Protease Inhibitor Cocktail Tablets. 10–30 mg of cell lysate was loaded in SDS-PAGE gels and transferred to polyvinylidene fluoride (PVDF) membranes. Membranes were blocked with Tris buffered saline-tween 20 (TBS-T) 5% non-fat milk buffer (w/v).

Primary antibodies are anti-mouse RLIM amino acids 1-271 (S691D third bleed; see MRC-PPU Reagents & Services http://mrcppureagents.dundee.ac.uk for further information and requests), anti-ERK1/2 (Santa Cruz Biotechnology) and anti-REX1 (Abcam). Secondary antibodies are Sheep IgG-horseradish peroxidase (HRP), Mouse IgG-HRP (Cell Signaling Technology) and Rabbit IgG-HRP (Cell Signaling Technology). After secondary antibody incubation, membranes were subjected to chemiluminescence detection with Immobilon Western Chemiluminescent HRP substrate (Millipore) using a Gel-Doc XR+ System (Bio-Rad) or to Infrared detection using a LI-COR Odyssey Clx system. Detected protein signals were quantified using Image J (NIH) or Image Studio (LI-COR Biosciences). All unprocessed immunoblots are provided in Supplementary Figure 2.

### Immunofluorescence

For localisation studies, mESCs were plated in 0.1% gelatin (v/v) coated coverslips and fixed in 4% PFA in PBS for 20 minutes at room temperature (RT). Cells were permeabilised with a 0.5% Triton X-100 in PBS solution for 5 minutes at RT and then blocked with 1% Fish gelatin (w/v) in PBS solution for 30 minutes at RT. RLIM primary antibody (Novus Biologicals) was diluted 1:200 in blocking solution and added to cells for 2 h at RT. Anti-mouse Alexa-488 (Thermo Fisher Scientific) was used as a secondary antibody at 1:500 in blocking solution for 1 h at RT. Actin Red 555 reagent (Thermo Fisher Scientific, one drop per ml of blocking solution) was added together with secondary antibody for actin staining. Hoescht was added at 1:10000 dilution in PBS for 5 min at RT as nuclear marker. Coverslips were mounted in glass slides using Fluorsave reagent (Millipore). Digital images were acquired in a Zeiss 710 confocal microscope and analysed and processed using ImageJ (NIH), Photoshop CC and Illustrator CC (Adobe).

### *In vitro* ubiquitylation assay

All recombinant proteins were produced in *E. coli* by MRC-PPU Reagents & Services and purified via standard protocols, and are available via http://mrcppureagents.dundee.ac.uk/. For *in vitro* ubiquitylation reactions, RLIM (140 nM) was incubated with a 20 μl ubiquitylation mix containing 0.1 μM UBE1, 0.05 μM UBE2D1 (UbcH5a) and 1.5 μg of REX1 (MRC-PPU Reagents & Services), 2 μM DyLightTM 800 Maleimide fluorescently labelled ubiquitin (Ub-IR^800^), 0.5 mM TCEP (pH 7.5), 5 mM ATP (both from Sigma Aldrich), 50 mM Tris-HCl (pH 7.5), 5 mM MgCl_2_ for 30 min at 30 °C. Reactions were stopped with SDS sample buffer and boiled for 5 min at 95 °C. Samples were loaded in 4-12% Bis-Tris gradient gels (Thermo Fisher Scientific). Gels were then scanned using an Odyssey CLx Infrared Imaging System (LICOR Biosciences). After scanning proteins were transferred to PVDF membranes and analysed via western blotting. All unprocessed gels and immunoblots are provided in Supplementary Figure 2.

### RNA extraction and quantitative RT-PCR

RNA was extracted using Omega total RNA extraction kit (column-based system) and obtained RNA converted to cDNA using iScript cDNA synthesis Kit (Bio-Rad). qPCR was performed using SsoFast EvaGreen Supermix (Bio-Rad) in a CFX384 real time PCR system (Bio-Rad). Relative mRNA levels were expressed using the ΔΔCT method and normalized to *Gapdh* expression. Data was analysed in Excel software and plotted using GraphPad Prism v7.0c software (GraphPad Software Inc.) (3). Primers used were: human *RLIM:* Forward (5’ to 3’): ATCATCAGGCTCATCAGGTGC, Reverse (3’ to 5’): AAGGAAGGGCAAAGAGCCAC; mouse *Xist*: Forward (5’ to 3’): GGATCCTGCTTGAACTACTGC, Reverse (3’ to 5’): CAGGCAATCCTTCTTCTTGAG: mouse *Gapdh:* Forward (5’ to 3’): CTCGTCCCGTAGACAAAA, Reverse (3’ to 5’): TGAATTTGCCGTGAGTGG. For *Xist* induction analysis, *Rlim*^+/y^ mESCs were cultured for 72 h in LIF-deficient mESC media prior to RNA extraction and analysis.

## DATA AVAILABILITY

The datasets generated during and/or analysed during the current study are available from the corresponding author on reasonable request.

## APPROVAL FOR HUMAN EXPERIMENTS

This patient was enrolled under the research study “Enhanced Care for Rare Genetic Diseases in Canada”. Research ethics approval for the study was obtained from the Children’s Hospital of Eastern Ontario Research Ethics Board (Ethics study number CTO 1577), which includes approval for sequencing and functional studies. All experiments were conducted in accordance within these ethical guidelines and regulations. Informed consent was obtained from all participants and/or their legal guardians.

## ACKNOWLEDGEMENTS

F.B. and G.M.F are supported by a Wellcome Trust/Royal Society Sir Henry Dale Fellowship (211209/Z/18/Z) and a Medical Research Council New Investigator Research Grant (MR/N000609/1). C.E-S. is supported by a Wellcome Trust PhD studentship and A.S-F. by a MRC-PPU prize studentship. Part of this work was performed under the Children’s Centre for Applied Translational Genomics Rare Diseases Functional Genomics programme and the Care4Rare Canada Consortium funded by Genome Canada and the Ontario Genomics Institute (OGI-147), the Canadian Institutes of Health Research, Ontario Research Fund, Genome Alberta, Genome British Columbia, Genome Quebec, and Children’s Hospital of Eastern Ontario Foundation. Part of this work was supported by the Luminesce Alliance – Innovation for Children’s Health, a not for profit cooperative joint venture between the Sydney Children’s Hospitals Network, the Children’s Medical Research Institute, and the Children’s Cancer Institute. It has been established with the support of the NSW Government to coordinate and integrate pediatric research. Luminesce Alliance is also affiliated with the University of Sydney and the University of New South Wales Sydney (A.J.E., K.D.K., M.J.W. and L.G.R). We thank Prof Miratul Muqit (MRC-PPU, University of Dundee) for critical reading of the manuscript.

## AUTHOR CONTRIBUTIONS

A.J.E. and K.D.K. provided exome analysis, M.J.W. provided clinical history, diagnosis and management of the patient and L.G.R. helped coordinate functional studies and prepared some figures. G.M.F. coordinated and F.B., C.E-S. and A.S-F. performed biochemical and cell-based experiments, analysed data and prepared figures. G.M.F. and L.G.R. wrote the paper with input from all authors.

## REFERENCES

1 Frints, S. G. M. et al. Pathogenic variants in E3 ubiquitin ligase RLIM/RNF12 lead to a syndromic X-linked intellectual disability and behavior disorder. Mol Psychiatry, doi:10.1038/s41380-018-0065-x (2018).

2 Hu, H. et al. X-exome sequencing of 405 unresolved families identifies seven novel intellectual disability genes. Mol Psychiatry 21, 133–148, doi:10.1038/mp.2014.193 (2016).

3 Tonne, E. et al. Syndromic X-linked intellectual disability segregating with a missense variant in RLIM. Eur J Hum Genet 23, 1652–1656, doi:10.1038/ejhg.2015.30 (2015).

4 Shin, J. et al. RLIM is dispensable for X-chromosome inactivation in the mouse embryonic epiblast. Nature 511, 86–89, doi:10.1038/nature13286 (2014).

5 Zhang, L. et al. RNF12 controls embryonic stem cell fate and morphogenesis in zebrafish embryos by targeting Smad7 for degradation. Mol Cell 46, 650–661, doi:10.1016/j.molcel.2012.04.003 (2012).

6 Bustos, F. et al. RNF12 X-Linked Intellectual Disability Mutations Disrupt E3 Ligase Activity and Neural Differentiation. Cell Rep 23, 1599–1611, doi:10.1016/j.celrep.2018.04.022 (2018).

7 Kumar, P., Henikoff, S. & Ng, P. C. Predicting the effects of coding non-synonymous variants on protein function using the SIFT algorithm. Nat Protoc 4, 1073–1081, doi:10.1038/nprot.2009.86 (2009).

8 Adzhubei, I., Jordan, D. M. & Sunyaev, S. R. Predicting functional effect of human missense mutations using PolyPhen-2. Curr Protoc Hum Genet Chapter 7, Unit7 20, doi:10.1002/0471142905.hg0720s76 (2013).

9 Schwarz, J. M., Rodelsperger, C., Schuelke, M. & Seelow, D. MutationTaster evaluates disease-causing potential of sequence alterations. Nat Methods 7, 575–576, doi:10.1038/nmeth0810-575 (2010).

10 Richards, S. et al. Standards and guidelines for the interpretation of sequence variants: a joint consensus recommendation of the American College of Medical Genetics and Genomics and the Association for Molecular Pathology. Genet Med 17, 405–424, doi:10.1038/gim.2015.30 (2015).

11 Gontan, C. et al. RNF12 initiates X-chromosome inactivation by targeting REX1 for degradation. Nature 485, 386–390, doi:10.1038/nature11070 (2012).

12 Shin, J. et al. Maternal Rnf12/RLIM is required for imprinted X-chromosome inactivation in mice. Nature 467, 977–981, doi:10.1038/nature09457 (2010).

13 Barakat, T. S. et al. RNF12 activates Xist and is essential for X chromosome inactivation. PLoS Genet 7, e1002001, doi:10.1371/journal.pgen.1002001 (2011).

14 Kernohan, K. D. et al. Diagnostic clarity of exome sequencing following negative comprehensive panel testing in the neonatal intensive care unit. Am J Med Genet A 176, 1688–1691, doi:10.1002/ajmg.a.38838 (2018).

