## Supplemental information for "A novel *RLIM/RNF12* variant disrupts protein stability and function to cause severe Tonne-Kalscheuer syndrome"

### Supplementary Figure 1

A Control

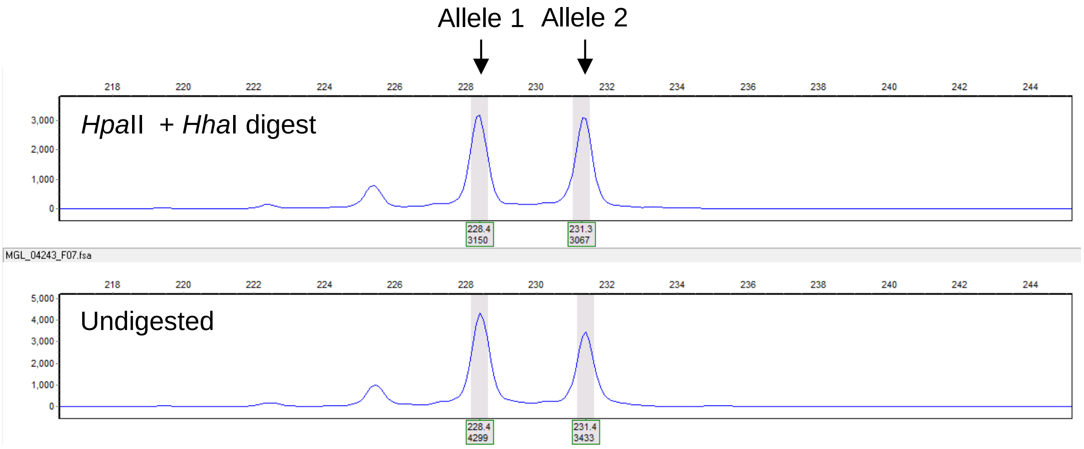


B II:2

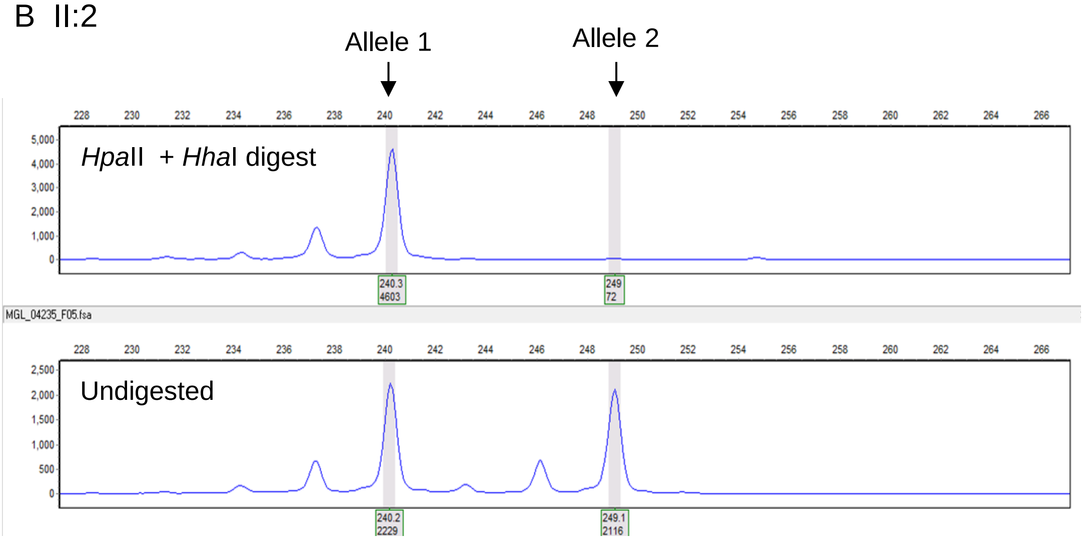


| Subject | Fragment length (bp) allele 1 | Fragment length (bp) allele 2 | Cells with allele 1 as active X-chromosome (%) | Cells with allele 2 as active X-chromosome (%) |
|---|---|---|---|---|
| Control | 228 | 231 | 50 | 50 |
| II:2 | 240 | 249 | 5 | 95 |

**Supplementary Figure 1:** Skewed X-inactivation in the proband's mother (II:2)**.** DNA was subjected to methylation sensitive restriction enzyme digestion, followed by PCR and fragment analysis. The inactive X-chromosome is not digested.

### Supplementary Figure 2: Unprocessed gels and blots

#### Figure 2A

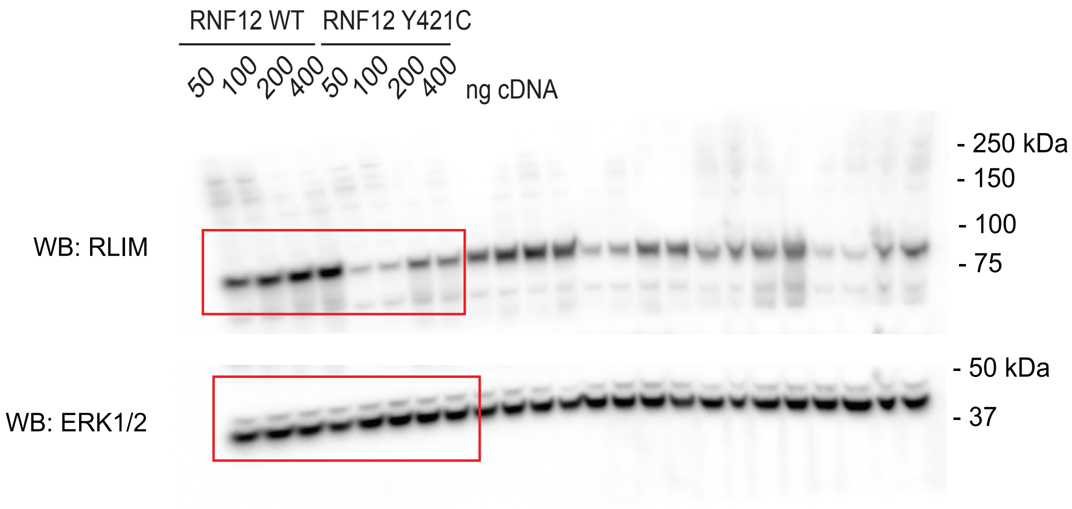


#### Figure 2C

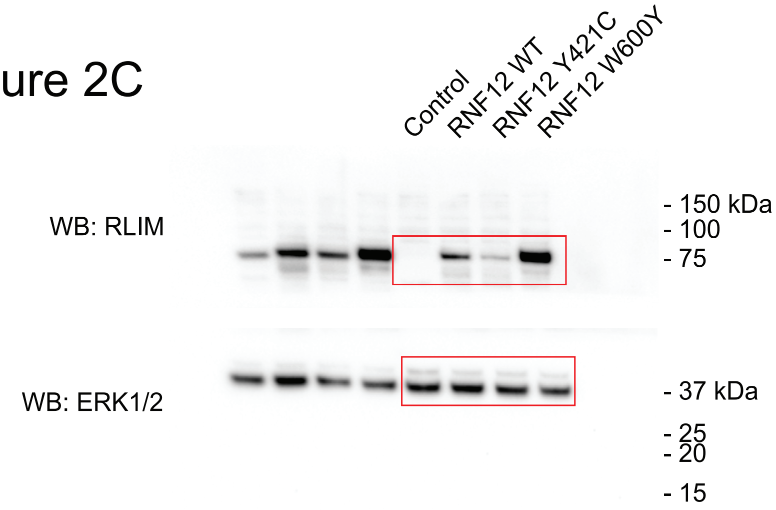


#### Figure 2E

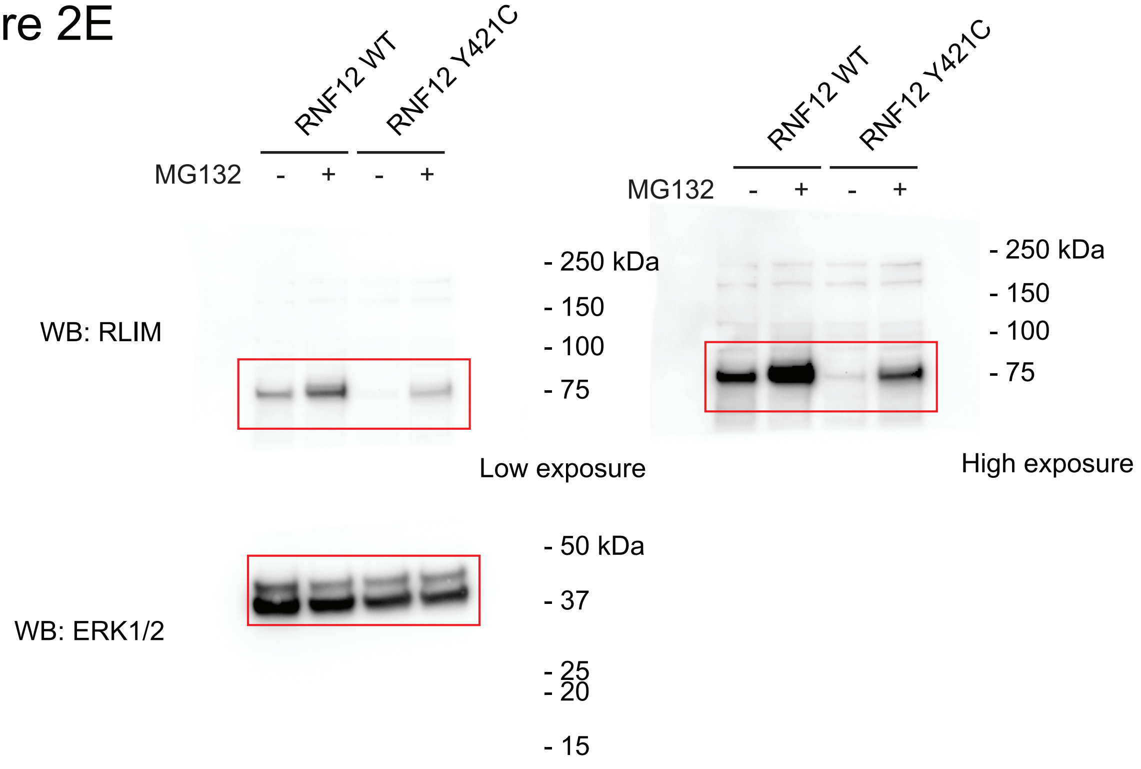

### Figure 3A

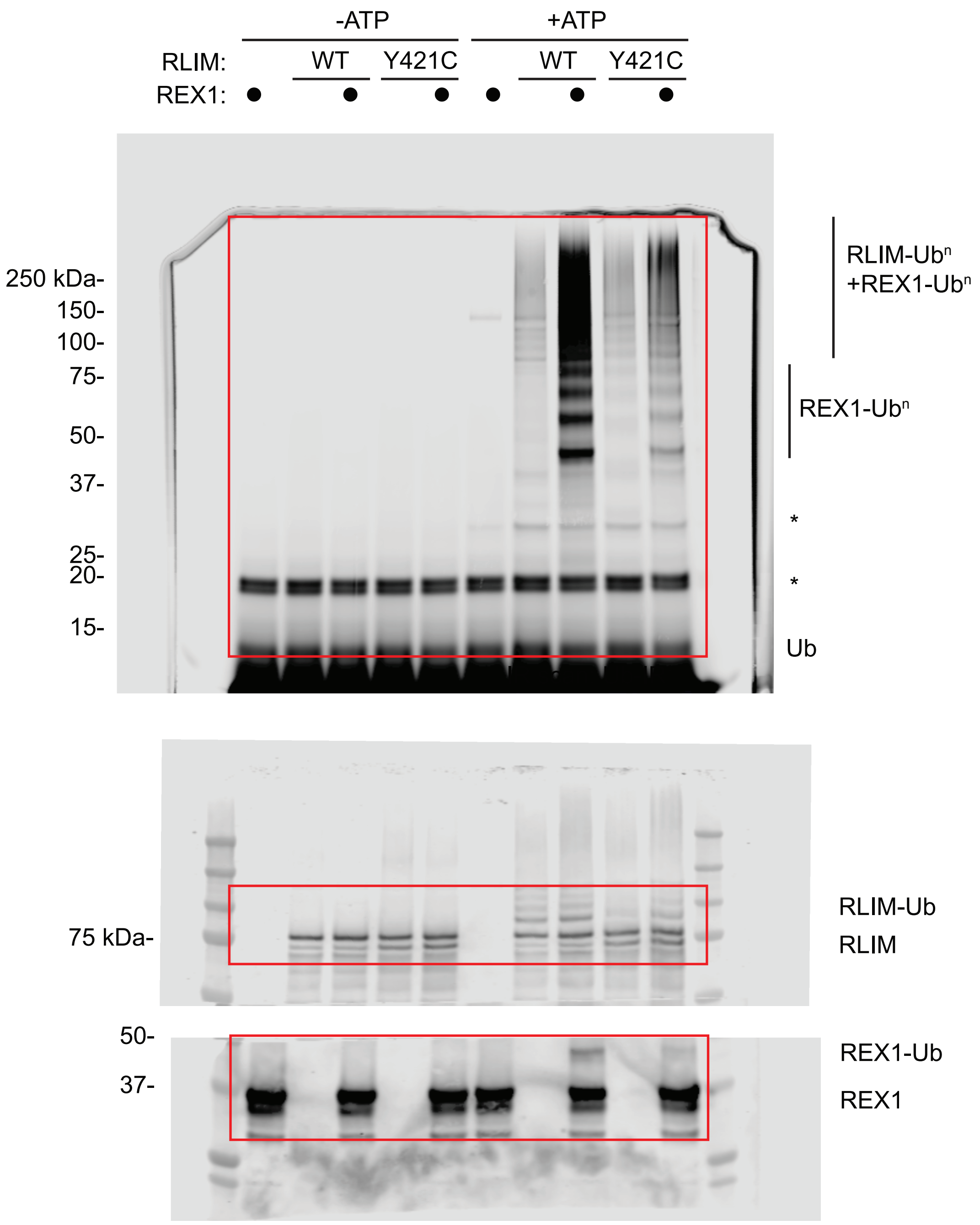


### Figure 4B

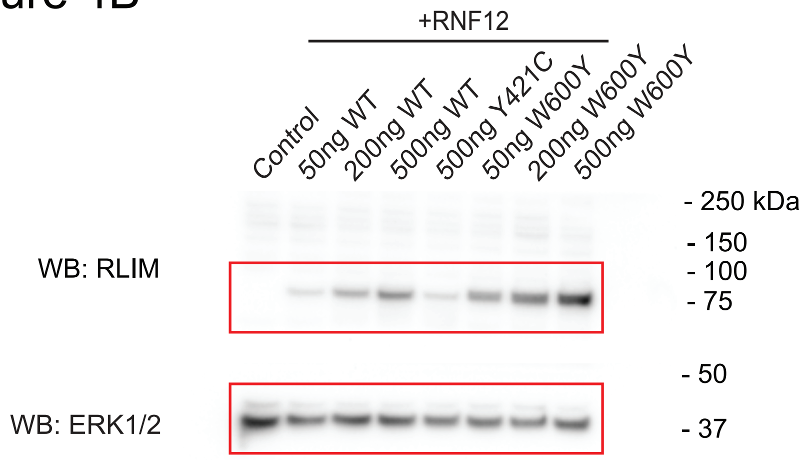
